# On the brain networks organization of individuals with high versus average fluid intelligence: a combined DTI and MEG study

**DOI:** 10.1101/2021.10.14.464389

**Authors:** S.E.P. Bruzzone, M. Lumaca, E. Brattico, P. Vuust, M.L Kringelbach, L. Bonetti

## Abstract

The neural underpinning of human fluid intelligence (G*f*) has gathered a large interest in the scientific community. Nonetheless, previous research did not provide a full understanding of such intriguing topic. Here, we studied the structural (from diffusion tensor imaging, DTI) and functional (from magnetoencephalography (MEG) resting state) connectivity in individuals with high versus average G*f* scores. Our findings showed greater values in the brain areas degree distribution and higher proportion of long-range anatomical connections for high versus average G*f*s. Further, the two groups presented different community structures, highlighting the structural and functional integration of the cingulate within frontal subnetworks of the brain in high G*f*s. These results were consistently observed for structural connectivity and functional connectivity of delta, theta and alpha. Notably, gamma presented an opposite pattern, showing more segregation and lower degree distribution and connectivity in high versus average G*f*s. Our study confirmed and expanded previous perspectives and knowledge on the “small-worldness” of the brain. Further, it complemented the widely investigated structural brain network of highly intelligent individuals with analyses on fast-scale functional networks in five frequency bands, highlighting key differences in the integration and segregation of information flow between slow and fast oscillations in groups with different G*f*.

## Introduction

A fundamental characteristic of the human brain is the ability to compute high-level logical, abstract reasoning and manipulate complex information to flexibly adapt to the environmental demands ^1–3^. This requires a set of cognitive skills, also referred to as fluid intelligence (G*f*), that are present across the population with measurable inter-individual differences ^1,3^. Indeed, G*f* refers to the ability of reasoning and solving logical and visual-spatial problems ^4,5^, involving a number of fine-grained cognitive abilities related to learning and memory. Due to their complex and fascinating nature, the investigation of G*f* and cognitive abilities have captured the attention of a large body of psychological and neuroscientific research ^6–14^, aiming to understand what is in the human brain that allows some individuals to outperform others in complex cognitive tasks. Nonetheless, the neural underpinning of individual differences in intellectual abilities is still far from being fully understood.

The human brain can be conceptualized as a complex system, whose efficiency arises from the balanced integration of activity coming from spatially segregated regions. In this framework, imbalances in brain network coordination have been linked to several psychiatric and neurological conditions ^15^ and modest network alterations have also been associated to the fine-grained differences in cognitive performances ^16,17^. Indeed, converging evidence suggests that human intelligence might depend on the organization of brain connectivity in a small-world network ^18,19^, a particular type of network where high connectivity between nodes is obtained with a relatively small number of connections ^9,20–23^, optimizing information flow across brain areas. This configuration implies a segregation of the network into independent, densely connected subnetworks (or modules) which are linked to other modules by a few, fundamental edges that allow to optimally integrate the information ^24^. Segregation properties of brain modules can be described by graph theory measures such as clustering coefficient and modularity. Conversely, we refer to integration as the property of the network to connect the non-overlapping modules through long-distance, crucial connections ^25^. In this case, cross-module functional integration properties can be described by characteristic path length, global efficiency, degree centrality and distribution and the presence of connector and provincial hubs ^26^. Thus, the key to fully understand the neural underpinning of fluid intelligence may relate to the configuration and flexibility of brain networks.

Nevertheless, classical studies on the neural basis of fluid intelligence provided evidence that has been organized within the framework of the Parieto-Frontal Integration Theory of intelligence (P-FIT) (Colom et al, 2010; Jung and Haier, 2007). According to this theory, cognitive performances arise from a chain of brain processes located in different regions such as occipital, temporal, parietal and frontal lobes. Indeed, incoming sensory information from temporal and occipital areas is first elaborated in parietal regions and subsequently integrated and abstracted in the frontal areas of the brain. The P-FIT theory is intriguing and coherent with several results described in years of research on intelligence. However, its approach tends to localise the main brain areas progressively involved in cognitive processes and did not directly considers the brain as a holistic dynamic system where integration and segregation are crucial to allow information flow and thus resolution of complex cognitive tasks. Along this line, other studies investigated the brain as a balanced network where integration and segregation of information play a crucial role. For instance, a growing body of evidence based on lesion ^27–29^ and functional magnetic resonance imaging (fMRI) studies pointed at a close link between fluid intelligence and a specific subset of brain regions that behave as brain hubs, which presumably mediate the information flow across different brain networks ^12,30,31^. This set of brain areas involves a widespread network comprising bilateral temporal, parietal and frontal regions, forming what is also referred to as “multiple demand” (MD) network ^12,30,31^. Furthermore, previous research studied the anatomical connectivity derived from fractional anisotropy (FA), a parameter commonly used to estimate the integrity of white matter tracts from diffusion tensor imaging (DTI) data ^32^. Studying such parameter is one of the main solutions to detect the strongest/weakest structural connections between brain areas as well as to estimate the network properties of the whole-brain. Remarkably, previous studies have associated enhanced FA in the superior longitudinal fasciculus, an association tract connecting frontal, parietal, temporal and occipital lobes, to greater scores in the Weschler Adult Scale of Intelligence (WAIS) for the fluid intelligence tasks ^8,33,34^. What is more, analysis of white matter network with graph theory reported higher global efficiency and shorter characteristic path length in participants with high versus average G*f* scores ^34,35^. Taken together, these studies suggested that the understanding of the neural underpinning of G*f* is progressively moving toward the network configuration of the whole-brain. However, the current available evidence did not return a clear picture of the brain organization of highly intelligent individuals. Further, there is not a full consensus about the most relevant properties of the brain networks to explain G*f*. On top of this, while previous works mainly focused on anatomical or functional connectivity using DTI and fMRI, evidence of functional connectivity based on electrophysiological methods is largely missing and needed. Indeed, although providing spatially accurate information, fMRI temporal resolution is extremely poor. In addition, it only provides an indirect measure of neural activity based on oxygen consumption and not on neuronal activity ^36–38^. In contrast, neurophysiological methods such as electroencephalography (EEG) and magnetoencephalography (MEG) detect direct brain activity with excellent temporal resolution, providing information at the milliseconds (ms) timescale ^39,40^. However, only a very limited number of studies explored the functional brain networks of G*f* using graph theory and EEG ^22,41^. The results of these studies pointed toward a small-world network configuration in individuals with greater G*f* scores and a main role of the parietal and frontal cortex for fluid intelligence, coherently with both the P-FIT and the MD network theories. Langer and colleagues ^22^ also reported that the clustering coefficient and characteristic path length of the functional brain network correlated to intelligence scores. Nonetheless, these studies relied on high-density EEG, and did not have an anatomical counterpart to confirm the results obtained from the neurophysiological results.

Thus, in this study we used MEG to explore the fine-grained differences in the brain networks of high versus average G*f* individuals as emerging from fast-scale whole-brain functional connectivity. Based on resting-state neural activity, we computed functional connectivity within five main frequency bands (delta: 0.1 – 2 Hz, theta: 2 – 8 Hz alpha: 8 – 12 Hz, beta: 12 – 32 Hz, gamma: 32 – 75 Hz) and investigated the properties of the emerging fast-scale networks with graph theory measures. Using the same measures, we explored the organization of the anatomical network and searched if the network (graph) properties of the two groups could be confirmed by microstructural changes in white matter.

## Results

### Experimental design and data analysis overview

In this study we aimed to characterize the neural correlates of fluid intelligence by using graph theory measures on functional and structural connectivity. To this goal, we acquired structural DTI using MRI and we measured brain activity with MEG during 10 minutes of resting state. Next, we collected behavioural measures of intelligence using the Wechsler Adult Intelligence Scale IV (WAIS-IV). The experimental procedures involved a total of 71 participants, but two participants had to be excluded since they did not perform the WAIS-IV tests. Our 69 WAIS-IV participants were divided into two groups based on their mean G*f* and by considering at least one standard deviation (std; standardized WAIS-IV std = 15) apart, so that the distinction between the two groups was psychometrically meaningful, as widely suggested by previous literature on the topic ^42^. The resulting groups were labelled as high G*f* (N = 38; mean G*f* = 117.72 ± 4.66) and average G*f* (N = 31; mean G*f* = 102.98 ± 6.09). As expected, the difference between the two groups was largely significant on a statistical level (t-test: *p* < 1.0e-07, *t*(55) = 11.08) (See Methods for further background and statistical information on the two groups). Finally, since we had to discard a few participants due to technical problems during the acquisition of DTI and MEG data, our final sample for WAIS-IV and DTI analysis consisted of 67 participants, while the one for the WAIS-IV and MEG analysis of 66 participants.

Back to the analysis, based on the non-cerebellar parcels of the automated anatomical labelling (AAL) brain parcellation, we constructed functional and structural connectivity matrices for each participant. The structural connectivity matrix was created based on the probabilistic tractography computed across all the 90 AAL regions of interest (ROIs) of the DTI images. The functional connectivity matrix was realized by reconstructing the sources of the neurophysiological signal acquired with MEG (using a beamforming algorithm) and by parcellating it with AAL. Importantly, the functional brain data was reconstructed in five different frequency bands (delta: 0.1 – 2 Hz, theta: 2 – 8 Hz alpha: 8 – 12 Hz, beta: 12 – 32 Hz, gamma: 32 – 75 Hz), returning a rather complete picture of the fast-scale information flow in the brain during resting state. Next, we computed graph theoretical measures of the individual brain structural and functional networks and compared them between the two groups of participants (high versus average G*f*).

Specifically, we were interested in the brain organization in terms of ROIs degree, segregation in different subnetworks (communities) and intra- and inter-subnetworks connectivity. Moreover, we aimed to detect how the brains of high versus average G*f* participants were organized in terms of structural connections and fast-scale information flow during resting state. Finally, we have complemented our network analysis with a comparison between high versus average G*f* groups in terms of white-matter tracts obtained computing tract-based spatial statistics (TBSS). The overview of the analysis pipeline is illustrated in **Figure 1**.

**Figure 1.**
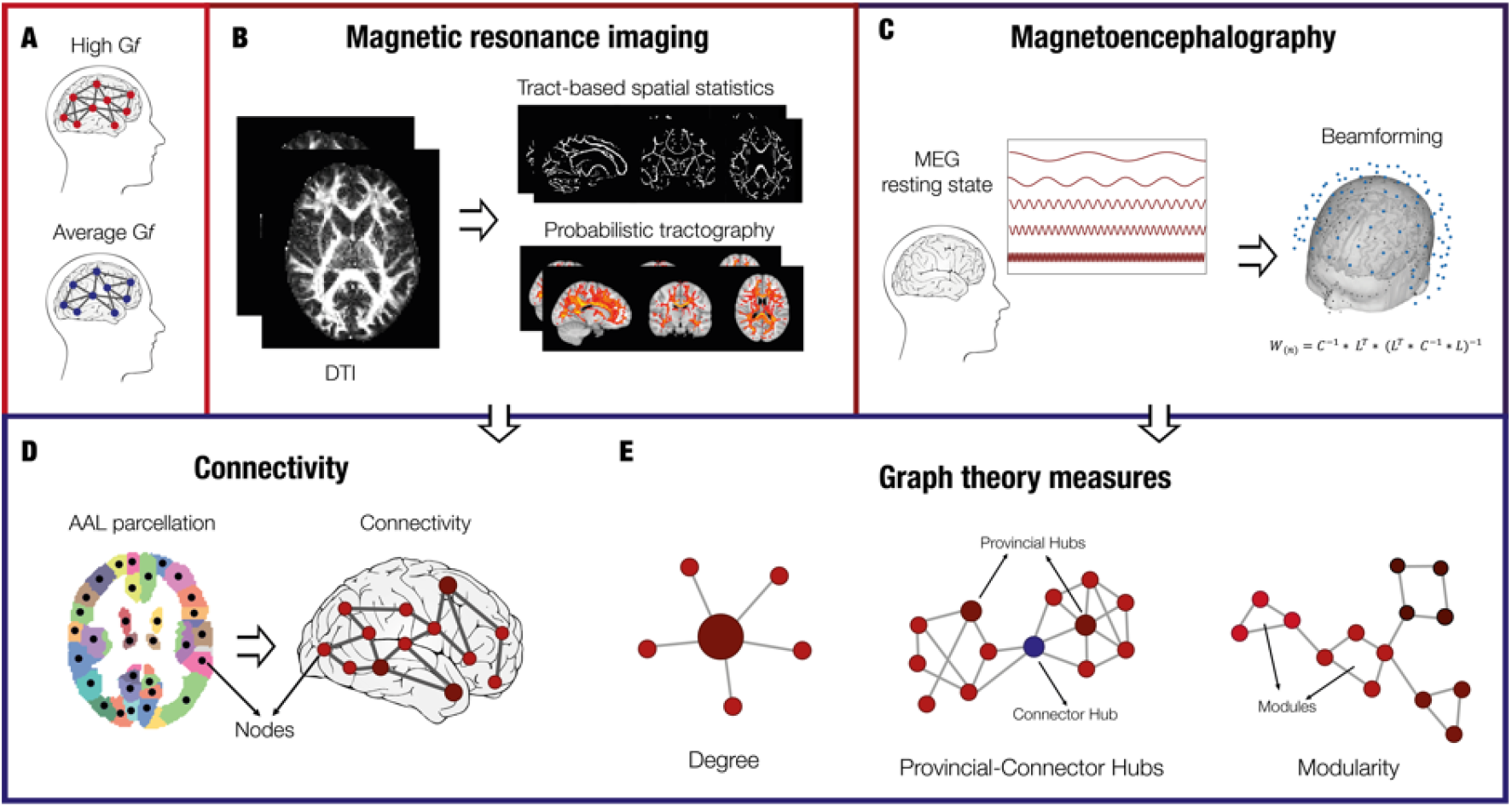
Experimental design and analysis pipeline. **A –** Participants were divided into two experimental groups, namely average Gf and high Gf, based on their scoring to perceptual reasoning, working memory and speed processing indexed by WAIS-IV. **B –** For both groups, diffusion-tensor imaging (DTI) data were collected and pre-processed. Then, differences in white matter microstructure were assessed with tract-based spatial statistics and the white matter bundles were modelled using probabilistic tractography. **C –** For both groups, magnetoencephalographic (MEG) data were collected during a 10-minute session of resting state. The data were filtered to analyse five different frequency bands: 0.1-2Hz (delta), 2-8Hz (theta), 8-12Hz (alpha), 12-32Hz (beta), 32-74Hz (gamma). Next, they were source-reconstructed with the beamforming algorithm. **D –** Connectivity was computed for both DTI and MEG data for each subject. The connectivity matrix for the DTI data was created by computing the probabilistic tractography based on AAL parcellation. The connectivity matrix for MEG data was estimated by computing the Pearson’s correlations between the envelope of each pair of brain areas. **E –**Graph measures were used to investigate the structural and functional brain networks of each group. Degree, provincial and connector hubs and modularity provided the most insightful results.

### Structural connectivity

After pre-processing the DTI data, matrices of structural connectivity were constructed for every participant using the output of the probabilistic tractography, which was normalized for the size of the brain ROIs (see Methods for details). We constrained the structural matrices to the non-cerebellar parcels of AAL parcellation (where each of the 90 regions represented a node of the brain network), resulting in a 90×90 matrix. The average structural connectivity across participants is showed in **Figure 2A**.

**Figure 2.**
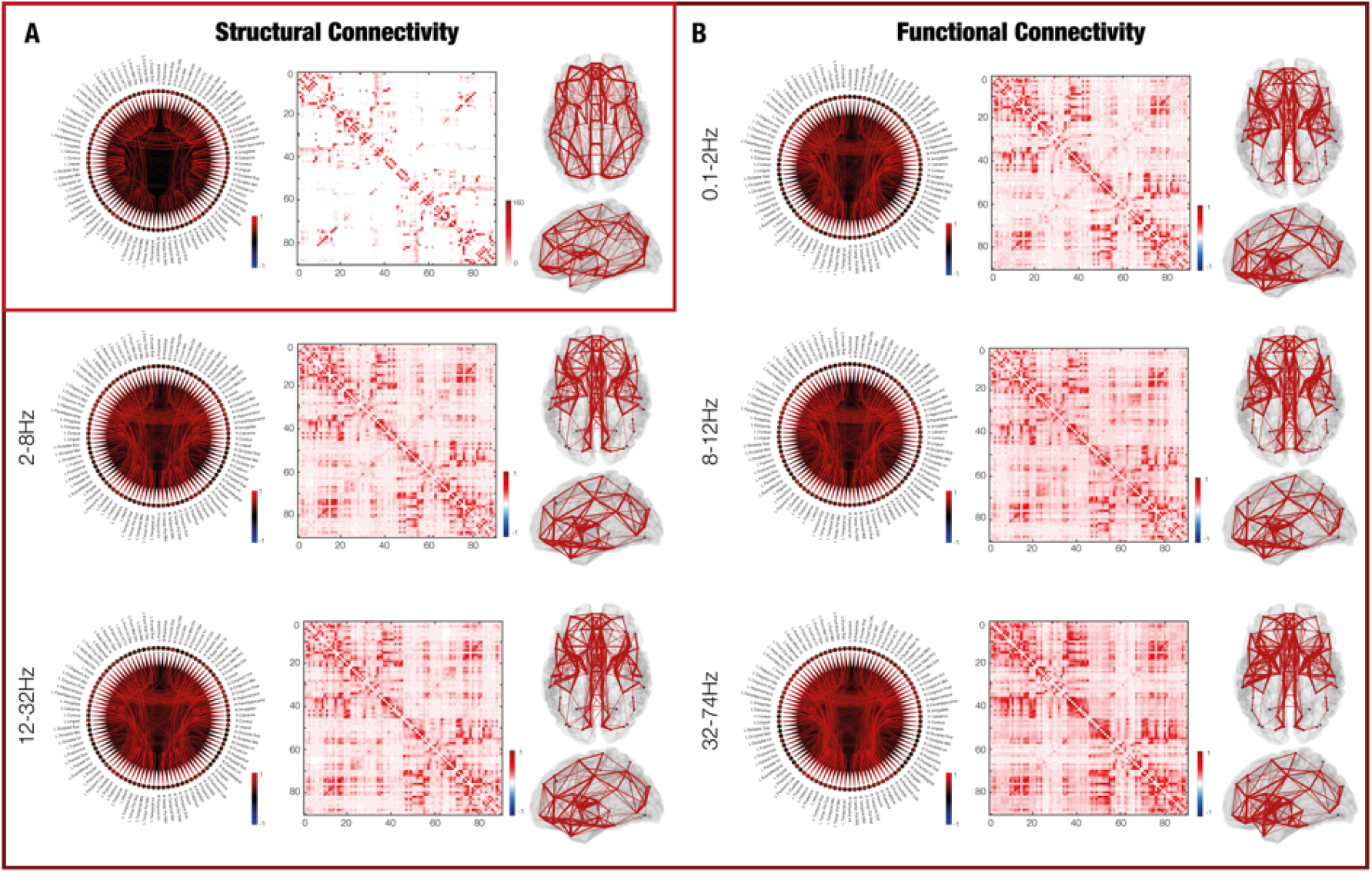
Structural and functional whole-brain connectivity. **A –** Structural connectivity computed from DTI data. The circular connectogram and the connectivity matrix represent the connections between the 90 AAL nodes. The different connection strengths are represented by different colour shades. The whole-brain figures depict the whole-brain connections, with stronger connections being thicker. Colourbars indicate the normalized average number of streamlines connecting the brain areas. **B –** Similarly, functional connectivity computed from MEG data, for each of the five frequency bands analysed. Colourbars indicate the Pearson’s correlations, showing the functional connectivity between brain areas.

**Figure 3.**
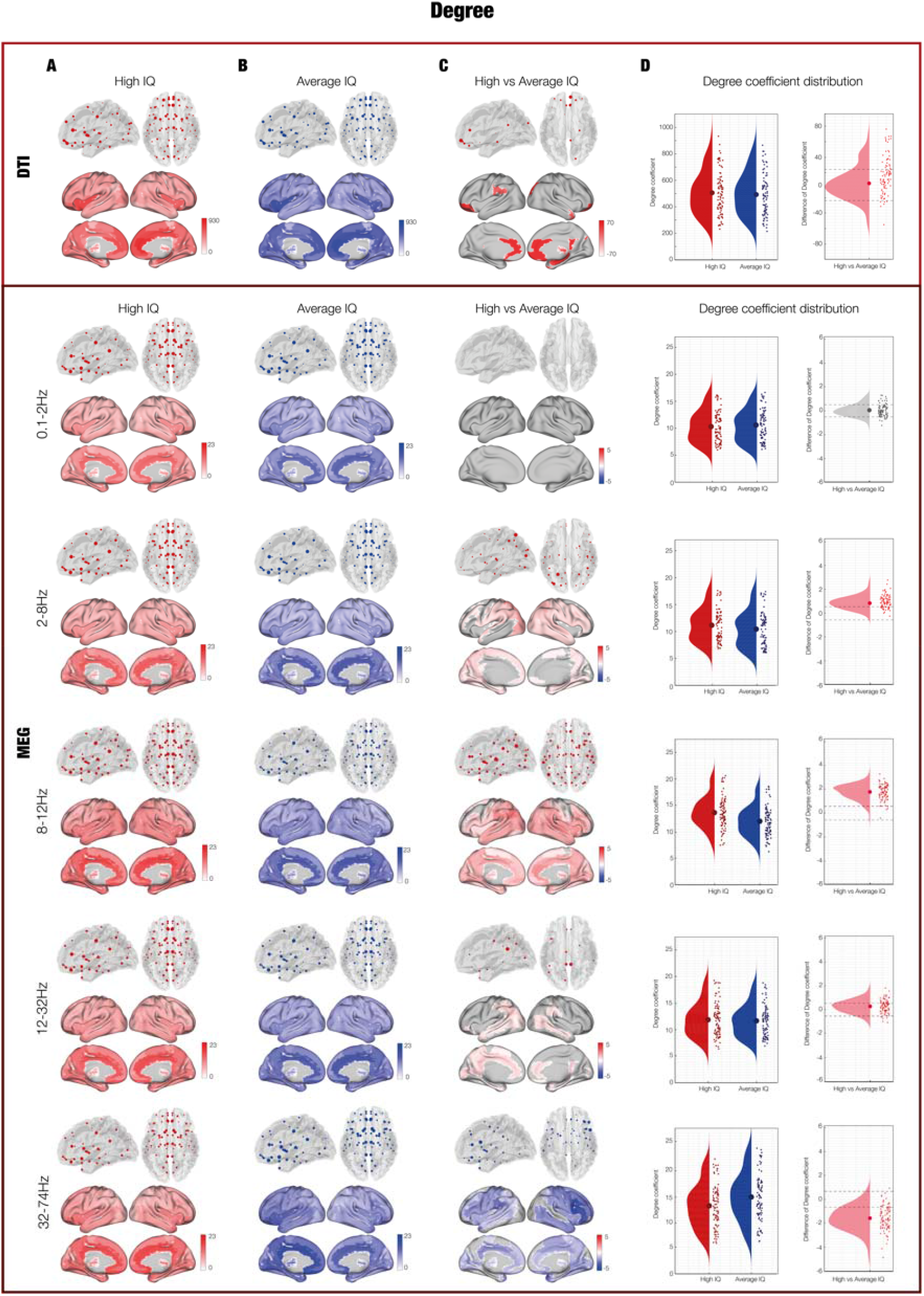
Degree of connectivity. **A –** Degree coefficients of structural and functional connectivity in participants with high Gf. **B –** Degree coefficients of structural and functional connectivity in participants with average Gf. **C –** Contrasts of the degree coefficients between the two groups. To be noted, high Gf individuals are represented in red, while average Gf ones in blue. In the contrast, the red colour indicates that high Gf individuals had a stronger degree distribution, while the blue showed a stronger distribution for average Gf participants. **D –** Degree coefficient distribution of high, average and high versus average Gf. Here, each dot shows the degree of each of the 90 ROIs, independently for high and average Gf participants. Dashed lines indicate the standard deviation with reference to zero, helping to identify how the degree distribution varied for high versus average Gf participants.

**Figure 4.**
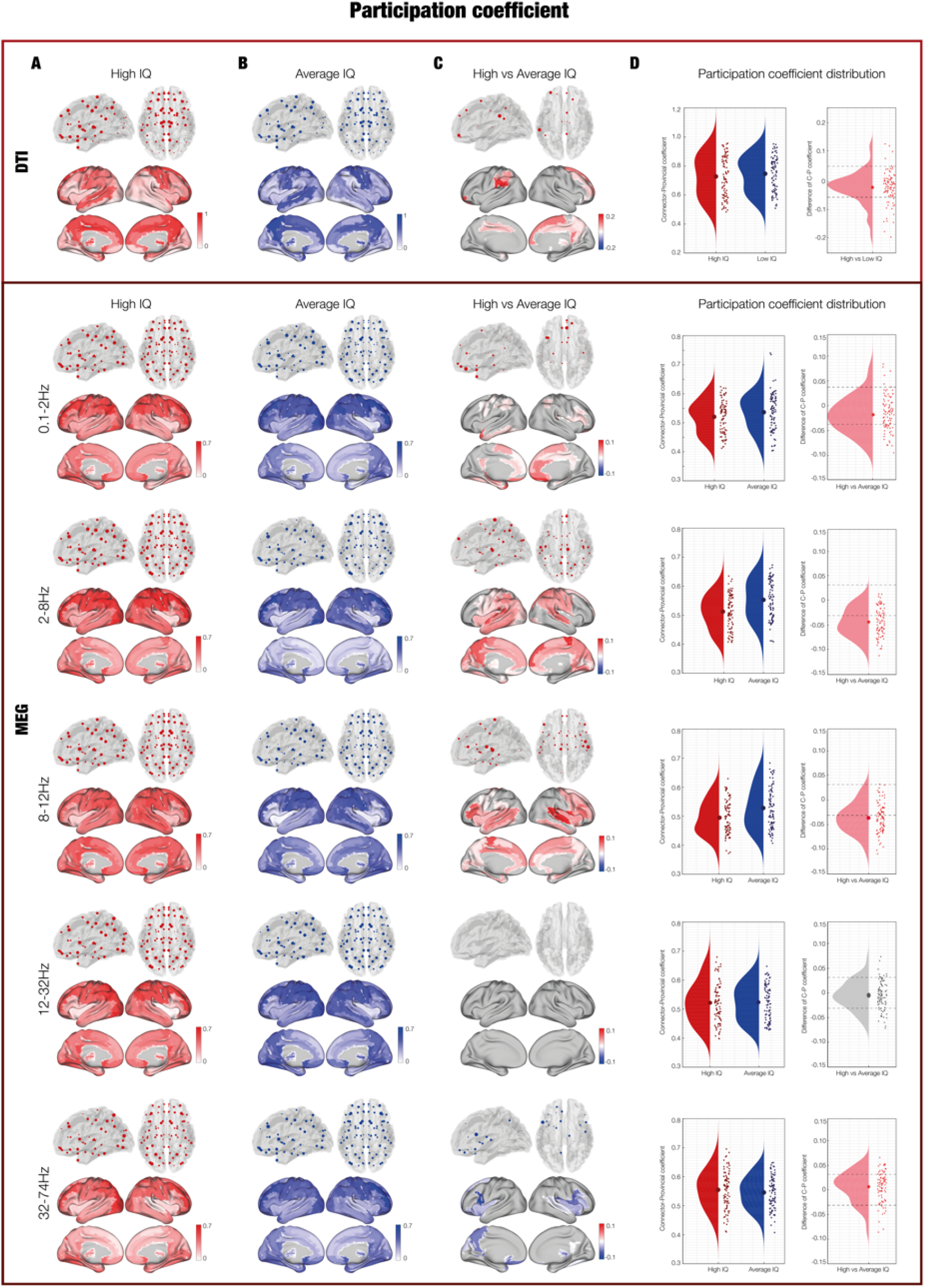
Connector and provincial hubs. **A –** Participation coefficient distribution computed from structural and functional connectivity in participants with high Gf. **B –** Participation coefficient distribution computed from structural and functional connectivity in participants with average Gf. **C –** Contrasts related to the Participation coefficient between the two groups. Positive results of the contrast indicate the presence of more intra- than inter-module in high versus average Gf participants. Conversely, the negative result of the contrast indicates more inter- than intra-module connections in high versus average Gf. To be noted, high Gf individuals are represented in red, while average Gf ones in blue. In the contrast, the red colour indicates that high Gf individuals had a more negative participation coefficient distribution, meaning that they presented a Participation coefficient more polarized toward the connector side. Conversely, the blue colour showed a stronger distribution of such coefficient for average versus high Gf participants. **d –** Participation coefficient distribution of high, average and high versus average Gf. Here, each dot shows the participation coefficient of each of the 90 ROIs, independently for high and average Gf participants. Dashed lines indicate the standard deviation with reference to zero, helping to identify how the participation coefficient distribution varied for high versus average Gf participants.

### Functional connectivity

Individual matrices of functional connectivity were constructed based on the pre-processed and source reconstructed MEG data, for each of the five frequency bands considered in the study: delta, theta, alpha, beta and gamma. As done for the DTI data, the reconstructed neural signal was constrained to the 90 non-cerebellar AAL parcellation. The resulting 90×90 matrix contained the information regarding the correlations between the 90 AAL brain regions, where each region represented a node of the brain network. The average functional connectivity across participants is shown in **Figure 2B**, independently for each frequency band.

### Graph theory measures

We analysed the two types of connectivity using graph theory measures between participants who scored high versus average in the WAIS-IV. For this purpose, we compared the various measures of the two groups with Monte Carlo simulations (MCS) to test the statistical significance.

#### Degree

First, we investigated whether the distribution of the ROIs degree was different among the two G*f* groups. Participants belonging to the high versus average G*f* group showed significantly higher distribution of degree in both structural (*p* = .007) and functional networks for theta (*p* < .001), alpha (*p* < .001) and beta (*p* = .004) frequencies, indicating an overall stronger level of connectivity between ROIs for the high G*f* participants. Remarkably, the main contributions to these values for structural connectivity and theta, alpha and beta frequency bands were provided bilaterally by a widespread network involving frontal (postcentral gyrus, superior frontal gyrus, postcentral gyrus, supplementary motor area), parietal (inferior and superior parietal lobule), occipital regions (inferior, middle and superior occipital gyrus) and temporal (middle and superior temporal gyrus) regions, as well as multiple subcortical areas (parahippocampal gyrus in the structural and in the functional, hippocampus, cingulum, thalamus in the functional). Conversely, individuals with average G*f* scores showed a greater degree distribution across the whole-brain compared to the high G*f* participants for the gamma frequency (*p* < .001). In this case, stronger degree centrality was observed in frontal, medio-temporal and subcortical areas, regions that greatly overlap to those that were more central for high versus average G*f* scores. A detailed list of the most central regions and the correspondent degree coefficients in structural and functional brain networks in the two experimental groups can be found in **Table ST1**. No significant difference was found for the distribution of degree in the delta frequency band.

#### Participation coefficient

First, we estimated the optimal community structure and modularity (depicted in **Figure 5** and reported in **Table ST2**) using the modularity algorithm introduced by Newman ^43^. Here, using MCS we tested the modularity values of structural and functional connectivity matrices (for the five frequency bands independently) against chance, to detect whether the brain networks were more modulable (more divisible into subgroups) than random configurations of the same original brain networks. The test was largely significant for both structural and functional connectivity matrices (*p* < .001).

**Figure 5.**
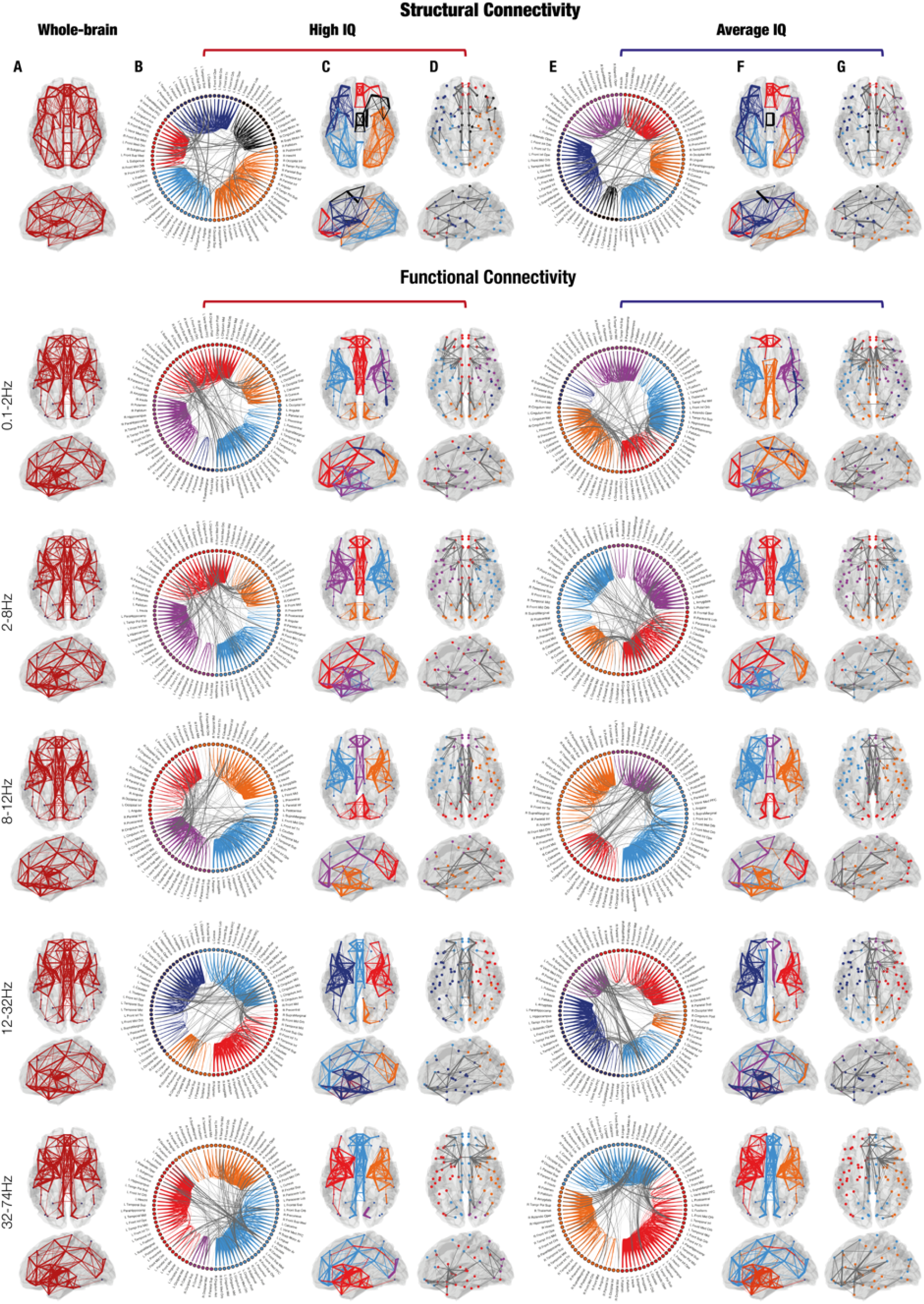
Inter and Intra-module connectivity in high versus average Gf. **A –** Whole-brain structural and functional connectivity in all participants. **B –** Circular connectogram representing inter-(in gray) and intra-module (different colors) connections in high Gf participants. **C –** Brain modules and intra-module connections overlaid on a standard brain template, in individuals with high Gf. Different modules are represented with different colors. **D –** Inter-module connections in individuals with high Gf. Different modules are represented by dots in different colors. **E –** Circular connectogram representing inter-(in gray) and intra-module (different colors) connections in average Gf participants. **F –** Brain modules and intra-module connections in individuals with average Gf. Different modules are represented with different colors. **G –** Inter-module connections in individuals with average Gf. Different modules are represented by dots in different colors.

Then, we computed the participation coefficient and compared it between the G*f* groups. This coefficient, ranging from zero to one, shows the level of connectivity of an ROI with the ROIs belonging to the same community when tending to one (provincial hub) or to ROIs of other communities when tending to zero (connector hub). Here, we studied the distribution of the participation coefficient in the high versus average G*f* participants. The results showed that high versus average G*f*s presented a higher distribution of connector than provincial hubs for both structural connectivity (*p* < .001) and delta (*p* < .001), theta (*p* < .001) and alpha (*p* < .001) bands of the functional networks. Main connector hubs for high G*f* individuals in these frequencies were found bilaterally in parietal, temporal, cingulate and subcortical areas (see **Table ST3**). Conversely, more provincial hubs (and thus more intra- than inter-community connections) were found in participants with average versus high G*f* for the gamma frequency band (*p* = .003) in frontal, temporal and subcortical regions (**Table ST3**). No differences were found between the two groups for the functional connectivity in the beta frequency band.

#### Modularity, Density, Characteristic path length, Global and Local efficiency

Modularity, density, characteristic path length, global and local efficiency were not significantly different between the two groups, neither in the structural nor in the functional networks. Nonetheless, before correcting for multiple comparisons, a significantly greater modularity distribution was found for the high versus average G*f* group in the theta (*p* = .046) and alpha (*p* = .029) frequency bands.

### Tract-based spatial statistics (TBSS)

Finally, to complement our network analyses, we performed TBSS to assess whether differential levels of fluid intelligence were associated to microstructural differences in white matter tracts.

The high versus average G*f* contrast revealed 38 clusters of significantly increased white matter, mostly located in frontal (postcentral gyrus, superior frontal gyrus, postcentral gyrus, supplementary motor area, precuneus, cingulum), temporal (temporal gyrus, parahippocampal gyrus,) and occipital (Calcarine fissure) regions. (see **Table ST1**). Instead, the average versus high Gf contrast revealed 32 significant clusters in analogous regions to those found in the first contrast (**Table ST2**) but with smaller dimensions, suggesting an overall modest, but noticeable, increase of white matter in high versus average G*f* individuals.

## Discussion

In this study, we investigated the fine-grained structural (DTI) and functional differences (MEG resting state) in individuals with high versus average G*f* scores. We found an overall increased degree in high compared to average G*f* individuals. Further, the two groups presented different community structures. Overall, in both structural and functional graphs the frontal brain areas of high G*f*s were grouped together in compact subnetworks including the cingulate gyrus and prefrontal regions. Conversely, frontal brain areas of average G*f*s belonged to more extended subnetworks which also included occipital and parietal regions. On top of these community structures, we have computed the participation coefficient, telling us if a brain area was principally connected to its own community or presented a high connectivity to external communities. Notably, brain areas of high versus average G*f*s were more connected to external communities, suggesting a stronger integration of brain subnetworks. Finally, microstructural analyses of white matter indicated a moderate but noticeable increase of white matter in high compared to average G*f* group.

### Intelligence and small-worldness of the brain

In this study, structural connectivity based on white matter tracts revealed an overall greater degree in the high versus average G*f* groups, as well as a greater trend of the brain areas to be connector instead of provincial hubs. This evidence suggests that the brain network configuration of individuals scoring the highest in G*f* tests is prevalently characterized by intermodular connections. Interestingly, although we computed analysis on the whole-brain distribution of degrees and participation coefficients, the brain regions that mainly contributed to such distribution were in frontal, hippocampal and cingulate areas, which were previously shown highly implicated in G*f* and cognitive processes ^12,16,33,44^. A further difference between the brain structural network of high and average G*f*s occurred in their community structure. Indeed, our modularity analyses grouped the frontal brain areas of high G*f*s together in compact subnetworks, while average G*f*s exhibited more extended subnetworks including at the same time frontal, parietal, and a few occipital regions. Notably, the medial cingulate gyrus of high G*f* belonged to a frontal subnetwork, while was segregated out from the rest of the brain areas in average G*f*. This finding suggests that the structural integration of the cingulate with frontal regions may be of key importance for fluid intelligence, coherent with several studies highlighting the role of the cingulate as a fundamental hub of the brain and its involvement in a plethora of cognitive processes ^33,44,45^.

In conclusion, the structural configuration of the brain of high G*f*s present properties of small-world networks, coherent with previous studies ^22,33–35,41^. Furthermore, such organization may provide the ideal wiring for efficient, long-range functional connections, and communication and integration between brain areas.

In line with this idea and differently from previous literature, we complemented our structural results with functional analyses of MEG resting state. Coherent with the structural organization that we have previously described, analyses of the functional graphs showed a higher functional degree across the whole-brain for three frequency bands: theta, alpha and beta. Additionally, we observed greater connector hub values in functional networks in delta, theta, and alpha power, reflecting the presence of more inter- than intra-module connections for the high G*f* group compared to the average G*f* group. In this case, the brain areas that mainly contributed to these results were sparser than in the structural graphs and belonged to temporal, frontal, parietal, and subcortical areas. Regarding the optimal community structure estimated for the two categories of participants, the results were compatible with the ones obtained for the structural graphs. Indeed, high G*f*s presented an overall stronger functional integration of the cingulate gyrus within frontal subnetworks of the brain, especially for delta, alpha, and beta. These findings suggest that better performance in fluid intelligence tasks is associated to an overall increased brain connectivity, which might reflect a more efficient signal integration favoured by a better inter-rather than intra-module communication between brain areas. Furthermore, the integration of the cingulate within frontal subnetworks of the brain suggests that the network organization of the cingulate may be of critical importance for individual differences in cognitive abilities and fluid intelligence.

Our results are overall coherent with previous literature. For instance, enhanced FA in long-range white matter tracts such as the superior longitudinal fasciculus have been associated to greater scores in intelligence tests ^8,34^. Further, measures of integration, segregation and “small-worldness” of the brain network have been associated to intelligence. Although we did not find significant differences for these measures in our cohort, higher global efficiency and shorter characteristic path length were found in participants with high versus average G*f* scores ^34,35^. Additional studies showed how modest network alterations were associated to the fine-grained differences in individual cognitive performances ^16,17^.

Taken together, previous works and our findings point at the idea that human intelligence depends on the organization of brain connectivity in a small-world network. Particularly, the key for understanding human intelligence may reside in the optimization of information flow across brain areas, which is made possible by balanced levels of integration and segregation, and short- and long-range connections in the human brain ^9,20,23^. In our study, on the one hand we further refined the understanding of such structural organization of the brain in relation to G*f*. On the other hand, for the first time we showed that the fast-scale functional networks detected with MEG presented similar features.

In contrast with previous studies ^34,35^, we did not find any significant difference regarding measures of modularity, characteristic path length, global and local efficiency. This might be due to the modest difference in the behavioural scores of G*f* in the two groups or may suggest that such broad whole-graph measures are not ideal to characterize the fine-grained essence of the brain of high-level performers in cognitive tests. Indeed, degree and participation coefficients might be more sensitive measures of integration and segregation network properties, being able to capture subtle but critical individual differences.

Finally, it is worth mentioning that microstructural analyses of white matter showed a moderate increase of white matter in the high compared to the average G*f* group. This increase in white matter might contribute to the greater scores in cognitive tests and the increased ability to integrate information across spatially distant brain areas in the high G*f* group. This finding would be coherent with studies showing a reduction of white matter integrity in ageing ^46^ and with declined cognitive abilities in clinical conditions ^47,48^.

### Network organization, intelligence and the role of cortical oscillations

While our results on the brain structural network organization confirmed and expanded previous literature, the finding on the brain functional networks provided rather new evidence that needs a deeper discussion. Indeed, our results showed a solid correspondence between brain structural and functional organization for what concerns delta, theta, alpha and beta frequency bands. In these cases, we observed very similar patterns of connectivity in high versus average G*f* individuals (e.g. enhanced degree and connector hub whole-brain distribution in high G*f* participants). Interestingly, we found an inverse pattern for the gamma frequency, where the average compared to the high G*f* group showed a greater degree distribution as well as a tendency of the brain nodes to be connector hubs.

Our results can be framed within the literature on cortical oscillations, which are thought to coordinate neuronal activity favouring communication across different brain structures ^49,50^. Specifically, delta (0.1-3 Hz) theta (4-8 Hz), and alpha (8-12 Hz) frequency bands have been repeatedly associated to complex cognitive functions. Delta and theta waves have been linked to response inhibition during attentional tasks ^51^ and to memory encoding and recognition tasks, respectively. Notably, the frequency peak for the theta frequency was found in the parietal cortex ^50,52^. Alpha frequency has been associated not only to sensory processes ^53^ but also to performance in academic results, memory and intelligence tests ^22,52^ In this respect, Langer and colleagues ^22^ reported an association between alpha power and psychometric intelligence scores, with strongest alpha activity in the right parietal cortex. Similarly, beta power has been reported to be positively associated to the strength of frontoparietal connectivity during visual search tasks ^54^. Although mostly based on task-related activity, previous evidence seems to point to a role for these frequency bands in cognitive functions contributing to fluid intelligence. In particular, the frequency peaks found in the frontal and parietal areas reported by these previous studies are coherent with the increased functional connectivity that we found in frontal, parietal, and temporal areas in our study. Spontaneously active brain regions are thought to reflect intrinsic properties of the brain, which in resting state may show the baseline functional organization of the information flow in the human brain. Notably, our results suggest that differences in terms of functional network organizations may represent a key to understanding human intelligence. Further, the nodes that we found to have significantly greater degree of connectivity and participation coefficient might well be important hubs for G*f* and consequently greater for high G*f* scores than for average ones. Thus, although our work points to a focus on integration and segregation of brain areas at the basis of fluid intelligence, our results are also coherent with the P-FIT theory, as well as the multiple demand network model for fluid intelligence that we have presented earlier.

Interestingly, gamma band presented a different behaviour when comparing high versus average G*f* people. Indeed, average compared to high G*f* had a greater degree distribution as well as a tendency of the brain nodes to be functional connector hubs. These results are in contrast with the other frequency bands and show that the functional resting state network in the gamma frequency presents more segregation and less information flow across the whole-brain in high versus average G*f*. Our results show differences between slower and faster cortical oscillation and may suggest that the most efficient brains (i.e. brains of more intelligent people) rely on gamma band for segregation of information within local subnetworks, while long-range functional communication and integration of information is mainly related to slower frequencies. Such evidence is coherent with previous literature which proposed gamma band for local communication of brain areas and slower frequencies such as alpha and theta for long-range functional connections ^55^. Nonetheless, future studies are called to replicate these results and expand our understanding of information flow across different frequency bands in relation to different levels of cognitive abilities and fluid intelligence.

## Conclusions

Altogether, our findings pointed to a different whole-brain configuration of connectivity between individuals scoring high versus average in G*f* tests. This was indicated by greater values in the brain areas degree distribution and by a higher proportion of long-range connections for the high versus average G*f* group. Further, the two groups presented different community structure, highlighting the structural and functional integration of the cingulate within frontal subnetworks of the brain in high versus average G*f*s. These results were consistently observed for structural connectivity and functional networks across slower frequency bands, especially delta, theta and alpha. Notably, only the faster frequency band, gamma, presented opposite results, showing more segregation and lower degree distribution and connectivity in high versus average G*f*s. In conclusion, this study confirmed and expanded previous perspectives and knowledge on the “small-worldness” of the brain. Further, it complemented the widely investigated structural brain network with analyses on fast-scale functional networks of five frequency bands, highlighting key differences in the integration and segregation of information flow between slow and fast oscillations.

## Methods

### Participants

We recruited a total of 71 healthy volunteers, 35 females and 36 males (aged 18-42, mean age: 25.06 ± 4.11 years) of different nationalities. Two participants had to be excluded since they did not perform the WAIS-IV tests. Further, for the DTI data (Tract-Based Spatial Statistics (TBSS) and the brain structural connectivity analyses), two participants were excluded from the sample due to the poor quality of the data, after the computation of the pre-processing pipeline. Thus, the final sample for DTI consisted of 67 healthy volunteers (34 females, 33 males, mean age: 24.94 ± 4.05 years). Regarding MEG, three participants were excluded because it was not possible to record their MEG resting state data. Thus, the final sample for the MEG functional connectivity analyses consisted of 66 healthy volunteers (34 females, 32 males, 24.95 ± 4.24 years).

All the experimental procedures were approved by the Ethics Committee of the Central Denmark Region (De Videnskabsetiske Komitéer for Region Midtjylland) (Ref 1-10-72-411-17), in compliance with the declaration of Helsinki – Ethical Principles for Medical Research.

### Experimental design and G*f* measures

Participants underwent the acquisition of functional (magnetoencephalography, MEG) and structural (magnetic resonance imaging, MRI) data. We recorded resting-state neurophysiological activity throughout 10 minutes of MEG recordings, during which participants were not engaged in any task and kept their eyes open. Regarding MRI, we acquired T1-anatomical and diffusion-weighted (DTI) brain images.

After acquiring the neuro-functional and -structural data, we collected behavioural measures to estimate the participants’ fluid intelligence measure (G*f*) along the following main scales of the fourth edition of the Wechsler Adult Intelligence Scale (WAIS-IV)^56^: perceptual reasoning, working memory and speed processing. All the tests were carried out in English, which was spoken fluently as a second language by the participants.

The mean G*f* score across the 69 (WAIS-IV subsample), 67 (WAIS-IV and DTI subsample) or 66 (WAIS-IV and MEG subsample) participants was nearly identical (111.10 ± 9.09; 111.45 ± 9.13 and 110.76 ± 9.05, respectively). Thus, the following numerical information about the two G*f* groups that we have used in our experiment will be reported for the full sample of 69 participants who were administered the WAIS-IV. Indeed, our sample was divided in two groups based on their mean G*f* and by considering at least one standard deviation (standardized WAIS-IV std = 15) apart, so that the distinction between the two groups was psychometrically meaningful, as widely suggested by previous literature on the topic ^42^. This procedure yielded two groups: the high G*f* group (N = 38; mean G*f* = 117.72 ± 4.66); the average G*f* group (N = 31; mean G*f* = 102.98 ± 6.09). As conceivable, the difference between the two groups was also statistically largely significant (*p* < 1.0e-07, *t*(55) = 11.08). Importantly, we controlled that the two groups were matched in terms of socio-economical, demographic, and educational status. In both groups, participants were mainly of Danish nationality and all of them came from a Western cultural country. The High G*f* group comprised 15 females and 23 males with an average age of 25.86 ± 4.89. The Average G*f* group comprised 18 females and 13 males with an average age of 24.00 ± 2.69. The age difference was not significant (*p* = .05). Furthermore, the mean of the education years was 14.73 ± 4.25 for the high G*f* and 14.56 ± 5.87 for the average G*f*. Neither this difference was significant (*p* = .37).

Back to the analysis pipeline, for each participant, we reconstructed the sources of the MEG signal by combining the MEG with the structural T1 MRI data in automated anatomical labelling (AAL) ^57^ space and estimated the functional connectivity between each pair of non-cerebellar brain areas of AAL. Similarly, we computed individual structural connectivity matrices based on the DTI images. Then, using graph theory measures, we analysed group differences for high versus average G*f* values in both structural and functional brain networks. The next paragraphs provide details about these procedures.

### Data acquisition

We acquired both MRI and MEG data at the Aarhus University Hospital (Denmark) in two independent sessions. MEG data were acquired with a 306-channel (204 planar gradiometers and 102 magnetometers) Elekta Neuroimag TRIUX system (Elekta Neuromag, Finland), with a sampling rate of 1000Hz and an analog filter of 0.1-330Hz. Prior to the measurements, the head shape and spatial coordinates of each participant were digitalizaed with a 3D digitizer (Polhemus FastrakColchester, VT - USA). The head localization was determined using four Head Position Indicator coils (cHPI) that were registered with respect to three anatomical landmarks (fiducials), namely the nasion, left and right preauricular areas. The cHPI allowed to continuously track the head position in respect to the MEG sensors and to correct for head movements. Furthermore, the digitalization of the participants’ head provided the information for co-registering the functional data recorded by the MEG with the anatomical data acquired with the MRI.

Whole-brain T1-weighted and diffusion-weighted images were acquired with a Siemens Magnetom Skyra 3T MRI scanner (20-channel head coils) located at Aarhus University Hospital, Denmark. T1 images were acquired with the following parameters: 1.0×1.0×1.0mm voxel size (1.0 mm^3^); 256×256 reconstructed matrix size; 2.96ms echo time (TE); 5000ms repetition time (TR); 240Hz/Px bandwidth. For the reconstruction of the MEG functional data, each T1-weighted scan was co-registered to the standard brain template from the Montreal Neurological Institute (MNI) using an affine transformation. Next, it was referenced to the MEG sensors space with the data about the head shape that was previously digitalized.

Diffusion-weighted images were acquired using echo-planar imaging (EPI), with the following parameters: 2.0×2.0×2.0mm voxel size (2.0mm^3^); 104ms TE; 3300ms TR; 100×100×72 matrix size; 221 volumes in anterior-posterior (AP) direction; 1 volume in posterior-anterior (PA) direction; 2500s/mm^2^ b-value; 29.41Hz/Px bandwidth.

### DTI data pre-processing

We pre-processed the MRI diffusion data with the FMRIB’s Diffusion Toolbox (FDT) toolbox in the FMRIB Software Library (FSL) ^58,59^. First, we visually checked the data to assess the good quality of the scans. After converting the files into *nifti* format, we created a reference volume (b0) based on the first image of both the AP and PA files, which we used to correct for susceptibility-induced distortions. Next, based on the corrected b0, we generated a brain mask that we applied to correct for head motion and eddy currents. The pre-processed and corrected data were subsequently used for the analysis of microstructural changes in white matter composition with Tract-Based Spatial Statistics (TBSS) and for the estimation of the main white matter tracts with probabilistic tractography.

### Tract-Based Spatial Statistics (TBSS)

We first computed the average fractional anisotropy (FA) images of each participant by fitting a tensor model to the pre-processed data. Next, we performed the analysis of microstructural changes in FA with TBSS ^60^, a series of functions of the FSL package that allow to compare the white FA values between groups of participants. The TBSS proceeds as follows. As a first step, likely outliers were removed by eliminating brain-edge artefacts and zeroing the end slices. Next, all the FA data were aligned into a common space, by means of a nonlinear registration performed on the FMRIB58_FA standard template (the FMRIB58_FA was obtained from a high-resolution average of FA images with 2×2×2mm spatial resolution, from 58 participants ^61^. Then, a mean skeleton representing the centers of all tracts common to the experimental group was created and taken into standard space (MNI152, 1×1×1mm). Finally, the skeletonized map of all participants was projected into a mean FA skeleton, with a threshold of 0.2. This procedure resulted in a final image representing the thickness of the white matter tracts independently for each participant. To compare such white matter tracts across the two G*f* groups, we computed t-tests for each white matter tracts voxel comparing values of high versus average G*f* participants. To correct for multiple comparisons, we adopted a cluster-based Monte-Carlo simulation (MCS) approach ^62,63^. This procedure assumes that the false positive results outputted by the t-tests would occur randomly and would therefore not be arranged in spatial clusters, while true significant results would form such clusters. Thus, in our MCS procedure, we have extracted the cluster of neighbouring significant voxels (where the difference between high versus average G*f* was significant), in the original data. Then, we computed 1000 permutations of the data. For each of the permutations, we have computed the clusters of significant values and extracted only the maximum cluster sizes. Such sizes gave rise to a reference distribution (built of the 1000 maximum cluster sizes extracted from the 1000 permutations) that we subsequently used to assess whether our original clusters were significant or not. Specifically, we have considered as significant the original clusters that were greater in size than 99.9% of the cluster sizes forming the permuted-based reference distribution.

### Tractography in AAL

We modelled the whole-brain structural connectivity with the FSL probabilistic tractography for crossing fibres ^64,65^, using the AAL parcellation. Based on the pre-processed data and the corrected reference volume b0, we estimated the fibre orientations of every voxel for each participant. Subsequently, we created 90 seed masks - one for each AAL region – with voxels sized 2×2×2mm. Using a Markov Chain Monte Carlo algorithm, we estimated the probability distribution of fibre direction at each brain voxel, with 1000 fibres (streamlines) per voxel. Whole-brain tracts (structural connectivity between each pair of AAL brain regions) were estimated by considering the continuity between fibres of all the voxels contained in each AAL region and all the other AAL regions.

### Structural connectivity network

After the estimation of the probabilistic tractography, we have computed a few normalization steps to obtain a final structural connectivity matrix, one for each participant.

In our brain networks, the nodes were defined according to the AAL parcellation, with each non-cerebellar AAL parcel representing a node of the network. The networks that we computed were undirected (i.e. a → b = b → a). However, the FSL probabilistic tractography estimates independently the two directions of the connectivity between two nodes (i.e. a → b = b → a means the same, but are estimated with slightly different values). Thus, as previously done ^15^, we averaged the two directions to obtain only one value of connectivity between any pair of brain areas and thus a truly symmetric undirected connectivity matrix. Finally, we have normalized each connection between AAL brain areas for the sizes of the same brain areas. This was done since larger AAL parcels may present more connections simply because they are larger and not because they are actually more densely connected. Thus, we have divided each connection between pairs of brain areas by the averaged size of those brain areas (e.g. a ← → b / ((size of a+ size of b)/2)). The resulting 90×90 matrix represented an undirected, weighted brain structural network.

### MEG data pre-processing

For the first pre-processing steps of the raw MEG data, we used MaxFilter ^66^. These steps consisted in applying signal space separation (SSS) to attenuate interferences originated outside the scalp, adjusting for head motion and down-sampling the signal from 1000Hz to 250Hz. Next, we converted the data into the Statistical Parametric Mapping (SPM) format and further proceeded with the analyses using the Oxford Centre for Human Brain Activity Software Library (OSL), a freely available toolbox that combines in-house-built functions with existing tools from FSL ^58^, SPM ^67^ and Fieldtrip ^68^ working in the Matlab environment (MathWorks, Natick, Massachusetts, United States of America). The frequencies below 0.1Hz, too low for being originated by brain activity, were removed with a high-pass filter. In addition, we applied a notch filter to correct for possible electric current-induced interferences and further down-sampled to 150Hz. After visually inspecting the data, we removed the parts of the signal that were altered by large artefacts. Then, we performed independent-component analysis (ICA) ^69^ to isolate and discard the artefacts generated by eyeblinks and heartbeat.

### Source reconstruction

The brain sources of the neural activity registered on the scalp by the MEG sensors were estimated by using the OSL implementation of the beamforming algorithm. Specifically, the forward solution was computed using an overlapping-spheres model in an 8-mm grid (comprising 3559 brain voxels). This solution represented a simplified geometric model of the MNI-co-registered anatomy of each participant, fitting a sphere separately for each MEG sensor ^70^. Then, we performed the inverse solution by using a beamforming algorithm. Such procedure utilized a different set of weights sequentially applied to the source locations for isolating the contribution of each source to the activity recorded by the MEG sensors at each time-point ^45,47^. Our beamforming computation was performed using both magnetometers and planar gradiometers.

Importantly, the source reconstruction was computed for five different frequency bands that were estimated after the ICA computation and subsequently reconstructed: delta: 0.1 – 2 Hz, theta: 2 – 8 Hz alpha: 8 – 12 Hz, beta: 12 – 32 Hz, gamma: 32 – 75 Hz.

### Functional connectivity network

After estimating the brain sources of the recorded MEG signal, we have computed one functional connectivity matrix for each participant, similarly to what we did for the structural connectivity based on the DTI data. First, the reconstructed functional data (3559 brain voxels) were constrained to the 90 non-cerebellar parcels defined by AAL. Next, we computed the envelope of the time-series from each brain region using the Hilbert transform. Finally, we estimated the functional connections between each pair of brain areas by computing Pearson’s correlations between the envelopes of the time-series of each pair of AAL brain regions.

### Graph theoretical measures

#### Degree of connectivity

The degree of connectivity describes how connected a node is to the other nodes of the network and can provide information about the functional integration properties of the network. We computed the degree (*d*_*(n)*_) of node *n* (here, an AAL ROI) as the sum of the strength of the connections of that node to all other nodes ^26^. This provided us with a value for each ROI indicating its degree of connectivity, and thus its centrality within the whole-brain network.

We tested the difference between high versus average G*f* participants using an MCS approach. Specifically, we computed the difference between the median of the degree of each ROI for high versus average G*f*. If the distribution of the degree is similar/equal between the two groups, their difference would be approximately zero, with some ROIs slightly above zero and some others slightly below, by random chance. Thus, in our MCS, we tested whether the distribution of differences between high versus average G*f* ROIs degree was significantly different from zero. First, we computed the number of ROIs whose difference degree was higher and lower than zero. Then we permuted the original data across experimental groups and computed the difference between the median of ROIs degree for the two permuted G*f* groups and observed the distribution of the difference between the degrees with respect to zeros. We re-iterated this operation for 10000 times, building a reference distribution of the difference between the ROIs degree in the permuted scenarios. Finally, we compared the original distribution of differences between high versus average G*f* ROIs degree with the permuted distribution. Since we tested the original distribution considering both tales of the permuted distribution (higher and lower than zero), the final MCS *p-value* was obtained by dividing the MCS α level by two (.05/2 = .025). Similarly, for the degree of functional connectivity, we performed 10 statistical tests: one for each of the two tales of the reference distributions and for each of the five frequency bands considered in the study. Thus, we corrected for multiple comparisons using the Bonferroni correction, by dividing the MCS α level (.05) by 10 (MCS *p-*value = .05/10 = .005).

#### Modularity and optimal community structure

Modularity is a value describing the optimal segregation of a network into discrete, non-overlapping clusters (modules) which optimize the network efficiency for specialized processing. In other words, it quantifies the degree to which a network can be subdivided into clearly defined, non-overlapping sub-networks. According to this definition, we computed the optimal community structure by maximizing the intra-module connections within non-overlapping sub-modules of the network and minimizing the inter-module connections. To calculate this measure, we used the undirected measure of modularity developed by Newman implemented in the Brain Connectivity Toolbox (BCT) (Rubinov and Sporns, 2010), relying on the eigenvector solution ^43^ and returning a discrete value of modularity and the corresponding optimal community structure, representing the division of the AAL ROIs into distinct, non-overlapping sub-networks of the brain.

We tested whether the modularity of the structural and functional brain data was significantly different by an equivalent network with connections placed randomly. To do so, we performed an MCS. First, we computed the modularity of the original data. Second, we performed 1000 permutations and extracted the modularity for the permuted data. This procedure yielded a reference distribution of permuted modularity values. Finally, we considered significant the original modularity value only if it was higher than the 99.9% of the permuted modularity values. This procedure was computed independently for the structural and functional data. A graphical depiction of the optimal community structure for structural and functional brain networks is provided in **Figure 5** and reported in detail in **Table ST2**.

#### Participation coefficient

Based on the previously computed optimal community structure, we were interested to observe whether the ROIs of high and average G*f* participants differed in terms of connectivity within and between the brain sub-networks. Specifically, we expected to find a tendency of high versus average G*f* individuals to have more pronounced connectivity between brain sub-networks. Thus, we computed the participation coefficient, which indicates whether an ROI is mainly connected to the other ROIs of the same sub-network or is more connected to ROIs in other sub-networks. The coefficient is computed by dividing the degree of the *ROI a* with regards to the ROIs of the same sub-network by the degree computed for *ROI a* with regards to all other ROIs (so also the ones of other sub-networks of the brain). Therefore, the coefficient values range between one and zero: the closer the coefficient to zero, the more the ROI has connections outside the community, highlighting its relevance as connector hub. Conversely, the closer the value to one, the greater the within-community degree, indicating that the ROI is mainly central within its own sub-network. To test the difference of the whole-brain distribution of the participation coefficient between high versus average G*f* individuals, we have performed an MCS analogous to the one described for the paragraph on the *Degree of connectivity*.

#### Characteristic path length

The *Characteristic path length* represents the average shortest path length between all pairs of nodes composing the network (e.g. the minimum number of connections to connect two nodes on average), providing a good estimate of how easily information flows through the network (and therefore of the integration of the network).

#### Global and Local efficiency

*Local efficiency* measures the average efficiency of integration within local clusters (e.g. between the neighbours of a given node). *Global efficiency* is the inverse of the characteristic path length and indicates how effectively the information flows across the network.

#### Density

*Density* represents the ratio between the number of actual edges of the network and the number of all possible edges of the network.

Each one of the measures described above (*characteristic path length, global and local efficiency, and density*) were statistically compared between high versus average G*f* groups by using two-sample t-tests. In this case, we corrected for multiple comparisons by using Bonferroni correction (i.e. dividing the α level of .05 by the total number of 24 comparisons (four measures x five frequency band of the functional networks plus one structural connectivity network), resulting in .05/24 = .002).

## Data availability

The codes are available at the following link: https://github.com/leonardob92/LBPD-1.0.git, while the multimodal neuroimaging data from the experiment are available upon reasonable request.

## Acknowledgements

We thank Giulia Donati, Riccardo Proietti, Giulio Carraturo, Mick Holt and Holger Friis for their assistance in the neuroscientific experiment. We also thank the psychologist Tina Birgitte Wisbech Carstensen for her help with the administration of psychological tests and questionnaires, and Francesco Carlomagno for his suggestions regarding the topic covered in the study.

The Center for Music in the Brain (MIB) is funded by the Danish National Research Foundation (project number DNRF117). Additionally, we thank the Italian section *of Mensa: The International High IQ Society* for the economic support provided to Francesco Carlomagno and the University of Bologna for the economic support provided to the students Giulia Donati, Riccardo Proietti, Giulio Carraturo.

LB is supported by the Carlsberg Foundation (grant CF20-0239) and by the Center for Music in the Brain, funded by the Danish National Research Foundation (DNRF117).

MLK is supported by the ERC Consolidator Grant: CAREGIVING (n. 615539), Center for Music in the Brain, funded by the Danish National Research Foundation (DNRF117), and Centre for Eudaimonia and Human Flourishing funded by the Pettit and Carlsberg Foundations.

## Author contributions

LB, SEPB, EB, MLK and PV conceived the hypotheses and designed the study. LB and SEPB performed pre-processing and statistical analysis. EB, MLK, ML, LB and PV provided essential help to interpret and frame the results within the neuroscientific literature. LB and SEPB wrote the first draft of the manuscript and prepared the figures. LB, SEPB, EB, ML, MLK and PV edited and reviewed the manuscript. All the authors contributed to and approved the final version of the manuscript.

## Supplementary Tables

The supplementary tables can be found at the following link: https://drive.google.com/drive/folders/1OSdaAbNoCO5zpQJrCYRKiHWoYgiEI882?usp=sharing

**Table ST1. Degree distribution**

Brain areas (ROIs) one standard deviation above (or below, as depicted by dash line in **Figure 3**) the mean degree. These ROIs provided the strongest contribution to the degree distribution that we tested statistically. The ROIs with the strongest values are the ones that had the highest difference in terms of degree when comparing High versus Average Gf groups (i.e. highest values correspond to ROIs that had a stronger degree for High versus Average Gfs). These areas are depicted in **Figure 3** in the brain templates. **Table ST1** reports ROIs independently for DTI and the five frequency bands from MEG.

**Table ST2. Community structure**

Brain areas (ROIs) reported in the different communities (modules) outputted by the modularity algorithm that we used in the study (Newman, 2006). The community structures are reported independently for DTI and the five frequency bands from MEG. These community structures are depicted in brain templates in **Figure 5**.

**Table ST3. Participation coefficient distribution**

Brain areas (ROIs) one standard deviation above (or below, as depicted by dash line in **Figure 4**) the mean participation coefficient. These ROIs provided the strongest contribution to the participation coefficient distribution that we tested statistically. In this case, the ROIs with the smallest values are the ones that had the highest difference in terms of participation coefficient when comparing High versus Average Gf groups (i.e. smallest values correspond to ROIs that had many more inter- than intra-community connections for High versus Average Gfs). These areas are depicted in **Figure 4** in the brain templates. **Table ST3** reports ROIs independently for DTI and the five frequency bands from MEG.

**Table ST4. TBSS – High versus Average Gf groups**

Significant clusters of stronger FA voxels when contrasting High versus Average Gf groups. The table reports Hemisphere, t-stat and MNI coordinates of each of the FA voxel forming the significant clusters.

**Table ST5. TBSS – Average versus High Gf groups**

Significant clusters of stronger FA voxels when contrasting Average versus High Gf groups. The table reports Hemisphere, t-stat and MNI coordinates of each of the FA voxel forming the significant clusters.

## References

1. Ashton, M. C., Lee, K., Vernon, P. A. & Jang, K. L. Fluid Intelligence, Crystallized Intelligence, and the Openness/Intellect Factor. J. Res. Pers. (2000). doi:10.1006/jrpe.1999.2276

2. Barbey, A. K., Koenigs, M. & Grafman, J. Dorsolateral prefrontal contributions to human working memory. Cortex (2013). doi:10.1016/j.cortex.2012.05.022

3. Gray, J. R., Chabris, C. F. & Braver, T. S. Neural mechanisms of general fluid intelligence. Nat. Neurosci. (2003). doi:10.1038/nn1014

4. Sternberg, R. J. Handbook of Intelligence. (Cambridge University Press, 2000).

5. Sternberg, R. J. The cambridge handbook of intelligence. The Cambridge Handbook of Intelligence (2019). doi:10.1017/9781108770422

6. Cattell, R. B. Theory of fluid and crystallized intelligence: A critical experiment. J. Educ. Psychol. (1963). doi:10.1037/h0046743

7. Kane, M. J. & Engle, R. W. The role of prefrontal cortex in working-memory capacity, executive attention, and general fluid intelligence: An individual-differences perspective. Psychon. Bull. Rev. (2002). doi:10.3758/BF03196323

8. Góngora, D., Vega-Hernández, M., Jahanshahi, M., Valdés-Sosa, P. A. & Bringas-Vega, M. L. Crystallized and fluid intelligence are predicted by microstructure of specific white-matter tracts. Hum. Brain Mapp. (2020). doi:10.1002/hbm.24848

9. Santarnecchi, E. et al. Network connectivity correlates of variability in fluid intelligence performance. Intelligence (2017). doi:10.1016/j.intell.2017.10.002

10. Criscuolo, A., Bonetti, L., Särkämö, T., Kliuchko, M. & Brattico, E. On the association between musical training, intelligence and executive functions in adulthood. Front. Psychol. 10, (2019).

11. Bonetti, L. et al. Auditory sensory memory and working memory skills: Association between frontal MMN and performance scores. Brain Res. 1700, 86–98 (2018).

12. Duncan, J., Assem, M. & Shashidhara, S. Integrated Intelligence from Distributed Brain Activity. Trends in Cognitive Sciences (2020). doi:10.1016/j.tics.2020.06.012

13. Bonetti, L. & Costa, M. Intelligence and Musical Mode Preference. Empir. Stud. Arts 34, (2016).

14. Bonetti, L. & Costa, M. Musical mode and visual-spatial cross-modal associations in infants and adults. Music. Sci. (2019). doi:10.1177/1029864917705001

15. Fernandes, H. M. et al. Disrupted brain structural connectivity in Pediatric Bipolar Disorder with psychosis. Sci. Rep. (2019). doi:10.1038/s41598-019-50093-4

16. Basten, U., Hilger, K. & Fiebach, C. J. Where smart brains are different: A quantitative meta-analysis of functional and structural brain imaging studies on intelligence. Intelligence (2015). doi:10.1016/j.intell.2015.04.009

17. Song, M. et al. Brain spontaneous functional connectivity and intelligence. Neuroimage (2008). doi:10.1016/j.neuroimage.2008.02.036

18. Bassett, D. S. et al. Cognitive fitness of cost-efficient brain functional networks. Proc. Natl. Acad. Sci. U. S. A. (2009). doi:10.1073/pnas.0903641106

19. Dwyer, D. B. et al. Large-scale brain network dynamics supporting adolescent cognitive control. J. Neurosci. (2014). doi:10.1523/JNEUROSCI.1634-14.2014

20. Buzsáki, G. Rhythms of the Brain. Rhythms of the Brain (2009). doi:10.1093/acprof:oso/9780195301069.001.0001

21. Srinivas, K. V., Jain, R., Saurav, S. & Sikdar, S. K. Small-world network topology of hippocampal neuronal network is lost, in an in vitro glutamate injury model of epilepsy. Eur. J. Neurosci. (2007). doi:10.1111/j.1460-9568.2007.05559.x

22. Langer, N. et al. Functional brain network efficiency predicts intelligence. Hum. Brain Mapp. (2012). doi:10.1002/hbm.21297

23. Eguíluz, V. M., Chialvo, D. R., Cecchi, G. A., Baliki, M. & Apkarian, A. V. Scale-free brain functional networks. Phys. Rev. Lett. (2005). doi:10.1103/PhysRevLett.94.018102

24. Deco, G., Tononi, G., Boly, M. & Kringelbach, M. L. Rethinking segregation and integration: Contributions of whole-brain modelling. Nature Reviews Neuroscience (2015). doi:10.1038/nrn3963

25. Zamora-López, G., Chen, Y., Deco, G., Kringelbach, M. L. & Zhou, C. Functional complexity emerging from anatomical constraints in the brain: The significance of network modularity and rich-clubs. Sci. Rep. (2016). doi:10.1038/srep38424

26. Rubinov, M. & Sporns, O. Complex network measures of brain connectivity: Uses and interpretations. Neuroimage (2010). doi:10.1016/j.neuroimage.2009.10.003

27. Duncan, J., Burgess, P. & Emslie, H. Fluid intelligence after frontal lobe lesions. Neuropsychologia (1995). doi:10.1016/0028-3932(94)00124-8

28. Roca, M. et al. Executive function and fluid intelligence after frontal lobe lesions. Brain (2010). doi:10.1093/brain/awp269

29. Woolgar, A. et al. Fluid intelligence loss linked to restricted regions of damage within frontal and parietal cortex. Proc. Natl. Acad. Sci. U. S. A. (2010). doi:10.1073/pnas.1007928107

30. Wen, T., Mitchell, D. J. & Duncan, J. Response of the multiple-demand network during simple stimulus discriminations. Neuroimage (2018). doi:10.1016/j.neuroimage.2018.05.019

31. Assem, M., Glasser, M. F., Van Essen, D. C. & Duncan, J. A Domain-general Cognitive Core defined in Multimodally Parcellated Human Cortex. bioRxiv (2019). doi:10.1101/517599

32. Basser, P. J. & Pierpaoli, C. Microstructural and physiological features of tissues elucidated by quantitative-diffusion-tensor MRI. J. Magn. Reson. (2011). doi:10.1016/j.jmr.2011.09.022

33. Hidese, S. et al. Correlation Between the Wechsler Adult Intelligence Scale-3rd Edition Metrics and Brain Structure in Healthy Individuals: A Whole-Brain Magnetic Resonance Imaging Study. Front. Hum. Neurosci. (2020). doi:10.3389/fnhum.2020.00211

34. Li, Y. et al. Brain anatomical network and intelligence. PLoS Comput. Biol. (2009). doi:10.1371/journal.pcbi.1000395

35. Van Den Heuvel, M. P., Stam, C. J., Kahn, R. S. & Hulshoff Pol, H. E. Efficiency of functional brain networks and intellectual performance. J. Neurosci. (2009). doi:10.1523/JNEUROSCI.1443-09.2009

36. Poldrack, R. A., Nichols, T. & Mumford, J. Handbook of Functional MRI Data Analysis. Handbook of Functional MRI Data Analysis (2011). doi:10.1017/cbo9780511895029

37. Berry, E. Chapter 2 -Image Processing Basics. A Pract. Approach to Med. Image Process. (2019).

38. Handbook of Functional Neuroimaging of Cognition. Handbook of Functional Neuroimaging of Cognition (2019). doi:10.7551/mitpress/3420.001.0001

39. Baillet, S. Electromagnetic brain mapping using MEG & EEG. Handbook of Social Neuroscience (2010). doi:10.1093/oxfordhb/9780195342161.013.0007

40. Gross, J. et al. Good practice for conducting and reporting MEG research. NeuroImage 65, 349–363 (2013).

41. Thatcher, R. W., Palmero-Soler, E., North, D. M. & Biver, C. J. Intelligence and EEG measures of information flow: Efficiency and homeostatic neuroplasticity. Sci. Rep. (2016). doi:10.1038/srep38890

42. Taylor, M. J. & Heaton, R. K. Sensitivity and specificity of WAIS-III/WMS-III domographically corrected factor scores in neuropsychological assessment. J. Int. Neuropsychol. Soc. (2001). doi:10.1017/s1355617701777107

43. Newman, M. E. J. Modularity and community structure in networks. Proc. Natl. Acad. Sci. U. S. A. (2006). doi:10.1073/pnas.0601602103

44. Langeslag, S. J. E. et al. Functional connectivity between parietal and frontal brain regions and intelligence in young children: The Generation R study. Hum. Brain Mapp. (2013). doi:10.1002/hbm.22143

45. Apps, M. A. J., Rushworth, M. F. S. & Chang, S. W. C. The Anterior Cingulate Gyrus and Social Cognition: Tracking the Motivation of Others. Neuron (2016). doi:10.1016/j.neuron.2016.04.018

46. Glahn, D. V. et al. Genetic basis of neurocognitive decline and reduced white-matter integrity in normal human brain aging. Proc. Natl. Acad. Sci. U. S. A. (2013). doi:10.1073/pnas.1313735110

47. Peng, X. et al. Reduced white matter integrity associated with cognitive deficits in patients with drug-naive first-episode schizophrenia revealed by diffusion tensor imaging. Am. J. Transl. Res. (2020).

48. Li, S. et al. Reduced integrity of right lateralized white matter in patients with primary insomnia: A diffusion-tensor imaging study. Radiology (2016). doi:10.1148/radiol.2016152038

49. Mahjoory, K., Schoffelen, J. M., Keitel, A. & Gross, J. The frequency gradient of human resting-state brain oscillations follows cortical hierarchies. Elife (2020). doi:10.7554/ELIFE.53715

50. Kahana, M. J. The cognitive correlates of human brain oscillations. Journal of Neuroscience (2006). doi:10.1523/JNEUROSCI.3737-05c.2006

51. Harmony, T. The functional significance of delta oscillations in cognitive processing. Frontiers in Integrative Neuroscience (2013). doi:10.3389/fnint.2013.00083

52. Klimesch, W. EEG alpha and theta oscillations reflect cognitive and memory performance: A review and analysis. Brain Research Reviews (1999). doi:10.1016/S0165-0173(98)00056-3

53. Başar, E. A review of alpha activity in integrative brain function: Fundamental physiology, sensory coding, cognition and pathology. International Journal of Psychophysiology (2012). doi:10.1016/j.ijpsycho.2012.07.002

54. Rogala, J., Kublik, E., Krauz, R. & Wróbel, A. Resting-state EEG activity predicts frontoparietal network reconfiguration and improved attentional performance. Sci. Rep. (2020). doi:10.1038/s41598-020-61866-7

55. Von Stein, A. & Sarnthein, J. Different frequencies for different scales of cortical integration: From local gamma to long range alpha/theta synchronization. in International Journal of Psychophysiology (2000). doi:10.1016/S0167-8760(00)00172-0

56. Wechsler, D. WAISLIII administration and scoring manual. The Psychological Corporation, San Antonio, TX (1997). doi:10.1177/1073191102009001003

57. Tzourio-Mazoyer, N. et al. Automated anatomical labeling of activations in SPM using a macroscopic anatomical parcellation of the MNI MRI single-subject brain. Neuroimage (2002). doi:10.1006/nimg.2001.0978

58. Smith, S. M. et al. Advances in functional and structural MR image analysis and implementation as FSL. in NeuroImage (2004). doi:10.1016/j.neuroimage.2004.07.051

59. Woolrich, M. W. et al. Bayesian analysis of neuroimaging data in FSL. Neuroimage (2009). doi:10.1016/j.neuroimage.2008.10.055

60. Smith, S. M. et al. Tract-based spatial statistics: Voxelwise analysis of multi-subject diffusion data. Neuroimage (2006). doi:10.1016/j.neuroimage.2006.02.024

61. Brown, C. A. et al. Development, validation and application of a new fornix template for studies of aging and preclinical Alzheimer’s disease. NeuroImage Clin. (2017). doi:10.1016/j.nicl.2016.11.024

62. Bonetti, L. et al. Rapid encoding of temporal sequences discovered in brain dynamics. bioRxiv (2020). doi:10.1101/2020.12.11.421669

63. Bonetti, L. et al. Spatiotemporal brain dynamics during recognition of the music of Johann Sebastian Bach. bioRxiv (2020). doi:10.1101/2020.06.23.165191

64. Behrens, T. E. J. et al. Characterization and Propagation of Uncertainty in Diffusion-Weighted MR Imaging. Magn. Reson. Med. (2003). doi:10.1002/mrm.10609

65. Jbabdi, S., Sotiropoulos, S. N., Savio, A. M., Graña, M. & Behrens, T. E. J. Model-based analysis of multishell diffusion MR data for tractography: How to get over fitting problems. Magn. Reson. Med. (2012). doi:10.1002/mrm.24204

66. Taulu, S. & Simola, J. Spatiotemporal signal space separation method for rejecting nearby interference in MEG measurements. Phys. Med. Biol. (2006). doi:10.1088/0031-9155/51/7/008

67. Penny, W., Friston, K., Ashburner, J., Kiebel, S. & Nichols, T. Statistical Parametric Mapping: The Analysis of Functional Brain Images. Statistical Parametric Mapping: The Analysis of Functional Brain Images (2007). doi:10.1016/B978-0-12-372560-8.X5000-1

68. Oostenveld, R., Fries, P., Maris, E. & Schoffelen, J. M. FieldTrip: Open source software for advanced analysis of MEG, EEG, and invasive electrophysiological data. Comput. Intell. Neurosci. (2011). doi:10.1155/2011/156869

69. Mantini, D. et al. A signal-processing pipeline for magnetoencephalography resting-state networks. Brain Connect. (2011). doi:10.1089/brain.2011.0001

70. Hillebrand, A. & Barnes, G. R. Beamformer Analysis of MEG Data. International Review of Neurobiology (2005). doi:10.1016/S0074-7742(05)68006-3

